# Proteomic Alterations Associated with Oral Contraceptive Use in Hypertensive Premenopausal African American Women

**DOI:** 10.1101/2024.08.12.607634

**Authors:** Lisa DeRoo, Malak Abbas, Gabriel Goodney, Merry L. Lindsey, Ashton F. Oliver, Han Le, Antwi-Boasiako Oteng, Amadou Gaye

**Author notes:** Corresponding author: Amadou Gaye, 1005 Dr DB Todd Jr Blvd, Nashville, TN 37208.

## Abstract

**Background:** Hypertension is a leading contributor to cardiovascular disease in reproductive-age women, with African American women experiencing disproportionate risk and severity. Oral contraceptive (OC) use has been implicated in elevating blood pressure, but the underlying molecular mechanisms remain unclear. Understanding the proteomic alterations associated with OC use may provide novel insights into the biological pathways contributing to OC-related hypertensive risk.

**Objective:** To identify proteomic signatures associated with oral contraceptive use among premenopausal African American women with stage 2 hypertension and elucidate pathways that may contribute to blood pressure regulation in this population.

**Study Design:** We evaluated 2,941 serum proteins measured on the Olink platform to assess associations with oral contraception use in 51 pre-menopausal women with untreated stage 2 hypertension (13 OC users and 38 non-users). A generalized linear model was fitted for each protein, adjusting for age, body mass index, alcohol consumption, level of education, and physical activity. Pathway enrichment analysis was carried out to identify common pathways among the proteins associated with oral contraceptive use. Gene Ontology enrichment analysis was conducted to gain insight into the functional characteristics of the proteins and the underlying biology.

**Results:** We identified 44 proteins significantly associated with OC use (FDR ≤ 0.05), including known and novel candidates. Among these, 13 proteins were elevated in OC users, including SERPINA6, SERPINA7, and TNFSF13, markers implicated in hormone transport, RAAS activation, and chronic inflammation. Proteins such as PLAU and PLAUR were enriched in sympathetic nervous system signaling and coagulation pathways, while SCG3 suggested neuroendocrine involvement. Enrichment analysis revealed overrepresentation of pathways related to complement and coagulation cascades, cholesterol metabolism, and the renin–angiotensin–aldosterone system..

**Conclusions:** OC use among premenopausal African American women, with stage 2 untreated hypertension, is associated with alterations in circulating proteins involved in neurohormonal signaling, immune modulation, lipid regulation, and vascular remodeling. The findings provide novel insights into the proteomic landscape associated with oral contraceptive in these women.

## INTRODUCTION

Hypertension remains a major public health burden and a key contributor to cardiovascular disease (CVD) mortality globally ^1^. Sex modulates both CVD risk and individual response to treatment, and understanding sex-specific differences in the pathophysiology of CVD is a necessary step for personalized medicine.^2^ In a nationally representative 2011-2016 survey conducted in non-pregnant women of reproductive age, 9.3% had hypertension; among those, 16.9% were unaware that they had hypertension and over 40% had uncontrolled hypertension.^3^ Cumulative exposure to elevated blood pressure (BP) over time is associated with increased risk of CVD outcomes and mortality, above and beyond current BP status. This suggests the importance of a life course approach to hypertension, with accumulated years of elevated BP in early to mid-adulthood having longer term implications for CVD risk.^4^ There are significant racial disparities for African Americans compared to Whites in prevalence, severity, and control of hypertension. African American women have a higher age-adjusted prevalence of hypertension (56.7%) than non-Hispanic White women (36.7%)^5^ and tend to develop hypertension at earlier stages of life and endure more severe manifestations of the condition.^6^ Among younger, premenopausal women with hypertension, African Americans were over two times more likely to have uncontrolled BP than whites. These disparities persist despite similar or even greater health awareness and healthcare engagement, underscoring the need to better understand context-specific risk factors in this population ^5^.

One such context is the use of OCs, a common hormonal method of birth control used by millions of women worldwide. Although generally safe, OC use has been associated with modest increases in BP and elevated CVD risk, particularly in women with pre-existing hypertension or additional risk factors such as smoking, obesity, or family history of CVD ^6-8^. Notably, women with stage 2 hypertension are often excluded from prospective trials of hormonal contraception, resulting in major evidence gaps regarding the biological effects of OCs in this vulnerable group ^9^.

Previous work has suggested that OC-associated hypertension may involve activation of the renin– angiotensin–aldosterone system (RAAS), increased sympathetic nervous system activity, and altered regulation of antidiuretic hormone (ADH) ^9-11^. Despite these known physiological effects, few studies have systematically profiled the circulating proteome to identify molecular changes linked to OC use in hypertensive women.

Among healthy, reproductive-aged women with a low absolute risk for CVD, the small increase in BP due to OC use is unlikely to be clinically relevant. However, for women with multiple risk factors (e.g., age > 35, high blood pressure), OC-use may increase CVD risk to an unacceptable level.

In this study, we sought to identify proteomic alterations associated with OC use in premenopausal African American women with stage 2 hypertension, a group at elevated risk for hypertensive complications yet underrepresented in biomedical research. Our aim is to (1) characterize the proteomic signature of OC use in this population and (2) explore the biological relevance of identified proteins in the context of known hypertension mechanisms.

## MATERIAL AND METHODS

### Study population

We used data from the GENomics, Environmental FactORs, and the Social DEterminants of Cardiovascular Disease in African Americans STudy (GENE-FORECAST), a study of 669 self-identified, U.S.-born African American men and women aged 21 to 65 years, recruited from the metropolitan Washington D.C. area during 2014-2022. The study protocol was approved by the Institutional Review Board of the U.S. National Institutes of Health (NIH), and written informed consent was provided by all participants. Data collection took place during a visit to the NIH Clinical Research Center. Pregnancy was an exclusion criterion for the study, and women were asked about menopausal status before undergoing a mandatory pregnancy test. Participants provided blood samples, underwent a physical exam and filled out questionnaires on medical history and health behaviors. BP measurements were taken in triplicate and averaged. OC use was obtained in a questionnaire aimed at collecting a comprehensive list of current medications.

GENE-FORECAST utilized a multi-omics systems biology approach, facilitating comprehensive, multi-dimensional characterization of health and disease among African Americans. Here, we isolated the differences in protein profiles for OC use among African American women with uncontrolled hypertension. Our inclusion criteria were women of pre-menopausal status with stage 2 hypertension (defined as systolic BP ≥ 140 mm HG or diastolic BP ≥ 90 mm HG based on the average of clinic measurements) who were not taking anti-hypertensive medication.

### Proteomic data

The ethylenediamine tetraacetic acid (EDTA) plasma samples from the GENE-FORECAST were sent to the Olink Proteomics Analysis Service in Boston, USA. All proteomic analyses were conducted collectively as one single batch. The Explore 3072 assay, utilizing eight Explore 384 Olink panels (Cardiometabolic, Cardiometabolic II, Inflammation, Inflammation II, Neurology, Neurology II, Oncology, Oncology II), were run to assess relative expression of a total of 2,947 unique proteins. Proximity Extension Assay echnology was conducted according to the Olink AB manufacturer procedures by the certified laboratory. Briefly, the technique relies on the use of antibodies labelled with unique DNA oligonucleotides that bind to their target protein present in the sample. The DNA oligonucleotides, when in proximity on a target protein, undergo hybridization and act as a template for DNA polymerase-dependent extension, forming a unique double-stranded DNA barcode proportionate to the initial protein concentration. Quantification of resulting DNA amplicons is accomplished through high-throughput DNA sequencing, generating a digital signal reflective of the number of DNA hybridization events corresponding to the protein concentration in the original sample. The measurement of protein levels is based on Normalized Protein eXpression (NPX) values, serving as a relative protein quantification unit. This quantification is normalized to account for systematic noise arising from sample processing and technical variation, leveraging internal controls and sample controls. NPX units are on a log2 scale, where a one NPX unit increase indicates a two-fold rise in the concentration of amplicons representing the target protein compared to the internal control. A total of 2,941 of the 2,947 proteins (99.8%) passed quality controls and were included in the analysis.

### Statistical analysis

Separate generalized linear models were fit for each of the 2,941 proteins, with protein level set as the outcome variable and main exposure of interest set as oral contraceptive use (no=0; yes=1), adjusting for age, body mass index (BMI), physical activity level, and education level. The p-values of the associations were adjusted for false discovery rate (FDR) using the Benjamini & Hochberg (BH) approach, with an FDR adjusted p-value ≤ 0.05 considered statistically significant.

Pathway enrichment analysis was undertaken to identify molecular pathways enriched in the set of proteins found to be associated with OC use. This analysis was carried out with the R algorithm *pathfinR*,^10^ a tool that defines subnetworks in a cluster of genes and identifies pathways over-represented in those subnetworks. Pathways with a BH adjusted enrichment p-value <0.05 were considered significantly overrepresented.

Gene Ontology (GO) enrichment analyses were subsequently conducted with the set of proteins associated with OC use to provide a systematic framework for annotating and deciphering biological functions, processes, and cellular components linked with the proteins and to gain insight into their roles within cellular and organismal contexts. GO analysis were conducted using the R libraries ‘enrichGO’ which runs a hypergeometric test to determine whether the set of proteins associated with OCs is enriched for specific GO terms compared to what would be expected by chance. GO terms with a BH adjusted enrichment p-value <0.05 were considered significantly overrepresented.

## RESULTS

Our study sample included 51 pre-menopausal women with stage 2 hypertension; 13 OC users and 38 non-users as after excluding 638 for reasons outlined in **Figure 1**. The 51 samples are decribed in **Table 1**. As expected, the two groups of OC users and non-users were similar in mean age (35 and 38 years, respectively) and mean BP (181 and 180 mmHg Systolic BP, respectively, and 72 mmHg Diastolic BP). Nearly all subjects were non-smokers. OC users were less likely to have a healthy BMI and more likely to drink alcohol, be a college graduate, and engage in moderate or greater than moderate physical activity than non-users.

**Table 1:** Baseline characteristics of the 51 women included in the analysis.

| Characteristics | HC (n = 13)<br>Mean/Count (SD %) | No HC (n = 38)<br>Mean/Count (SD %) | P-Value |
| --- | --- | --- | --- |
| **Age (years)** | 35 (10) | 38 (9) | 0.30 |
| **Systolic Blood Pressure (mmHg)** | 181 (20) | 180 (23) | 0.94 |
| **Diastolic Blood Pressure (mmHg)** | 72 (13) | 72 (10) | 0.93 |
| **Body Mass Index (kg/m<sup>2</sup>)** | 32 (6) | 31 (8) | 0.79 |
| **Current Smoker** |  |  |  |
| No | 13 (100%) | 36 (95%) | 1.00 |
| Yes | 0 (0%) | 1 (3%) |  |
| **Drinks Alcohol** |  |  |  |
| No | 2 (15%) | 12 (32%) | 0.41 |
| Yes | 11 (85%) | 25 (66%) |  |
| **Education Level** |  |  |  |
| ≤ High School | 2 (15%) | 11 (29%) | 0.38 |
| High School + Vocational Educational | 4 (31%) | 13 (34%) |  |
| ≥ Graduate Degree | 7 (54%) | 12 (32%) |  |
| **Physical Activity (Baeckes' sport index)** |  |  |  |
| < Moderate Activity | 5 (38%) | 17 (45%) | 0.94 |
| ≥ Moderate Activity | 8 (62%) | 21 (55%) |  |

**Figure 1:**
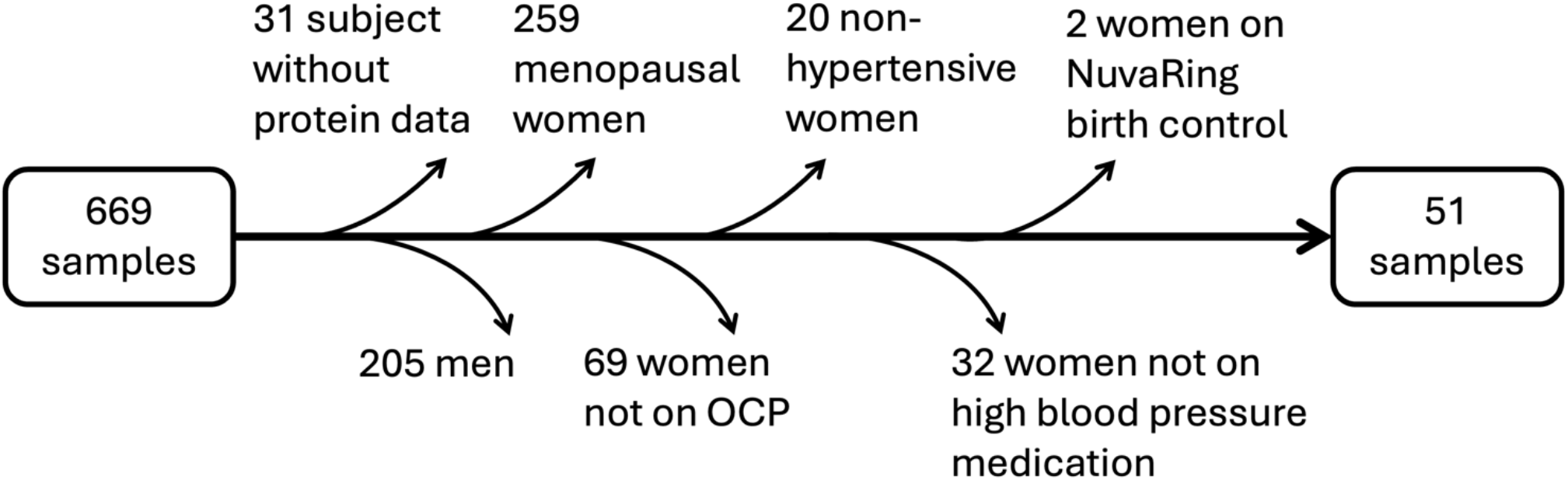
Flowchart illustrating sample selection: of 669 pre-menopausal women screened, 618 were excluded based on criteria detailed in the diagram, resulting in a final analytic sample of 51 women.

**Figure 2:**
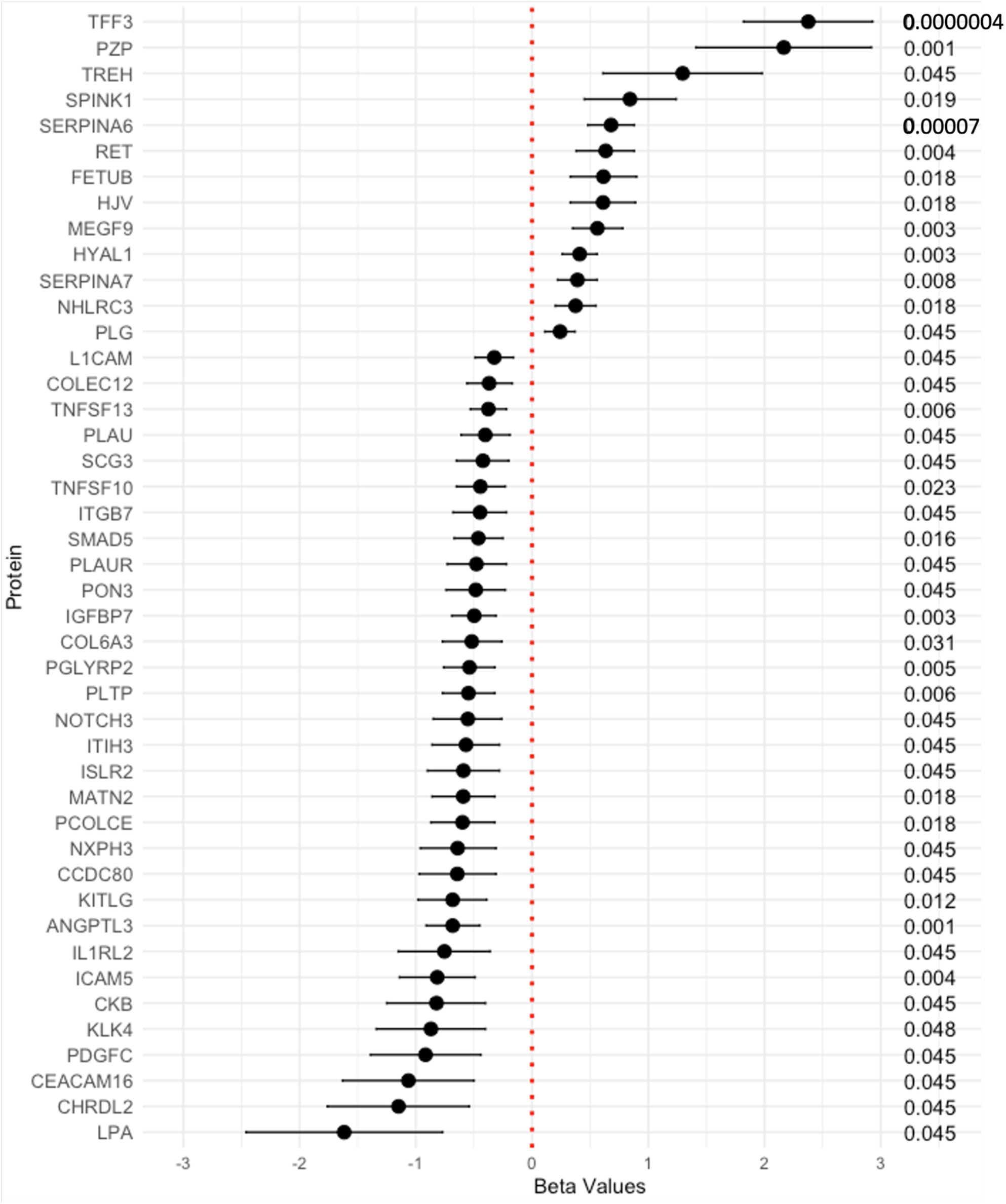
Proteins linked to hormonal contraceptive use, with beta coefficients and corresponding 95% confidence intervals and adjusted p-values. The genes are listed from negative to positive beta coefficient.

A total of 2,941 (99.8%) out of 2,947 proteins passed quality control. All the measurements were conducted in a single, centralized batch ensuring minimal batch effects and high internal consistency. The technology employed by Olink offers high specificity and sensitivity by relying on dual antibody recognition and DNA barcoding, reducing cross-reactivity and improving signal fidelity. Built-in internal and external controls enabled stringent normalization and correction for technical variation.

### Proteins Associated with OC Use

We identified 44 proteins significantly associated with oral contraceptive use (Figure 1 and Supplemental Table T1). There were 31 proteins exhibiting lower levels among OC users and 13 proteins with higher levels. We identified 44 proteins significantly associated with OC use; the results are summarized graphically in Figure 1 an detailed in Supplemental Table T1. Of these 44 proteins, 31 were found at lower levels and 13 at higher levels among OC users. Notably, several proteins previously implicated in blood pressure regulation and hormonal response pathways were differentially expressed.

Elevated levels of SERPINA6 and SERPINA7, key hormone-binding proteins, point toward activation of the renin–angiotensin–aldosterone system (RAAS). Similarly, higher expression of TNFSF13, involved in inflammatory signaling, and SCG3, a neuroendocrine granin protein, supports a role for inflammation and sympathetic nervous system activation in OC-associated hypertension. We also observed increases in PLAU and PLAUR, which contribute to both coagulation and neurovascular signaling.

Conversely, proteins such as ANGPTL3 and NOTCH3, involved in lipid metabolism and vascular remodeling respectively, were downregulated in OC users, suggesting additional pathways of vascular and metabolic dysregulation. Novel findings include differential expression of SMAD5, potentially linking OC use to TGF-β/BMP-mediated vascular remodeling.

### Pathway Enrichment Analysis

Pathway enrichment analysis of the 44 differentially expressed proteins revealed multiple significantly enriched biological pathways (FDR ≤ 0.05), outlined in **Figure 3**. The most significant pathway was complement and coagulation cascades (adjusted p = 2×10^−5^), supported by the involvement of PLAU, PLAUR, and PLG, which are key regulators of fibrinolysis and vascular homeostasis. Additional enriched pathways included prostate cancer, proteoglycans in cancer, and cholesterol metabolism, involving proteins like ANGPTL3 and LPA. Immune and metabolic signaling pathways were also highlighted, including TGF-beta signaling (SMAD5), Notch signaling (NOTCH3), and PPAR signaling.

**Figure 3:**
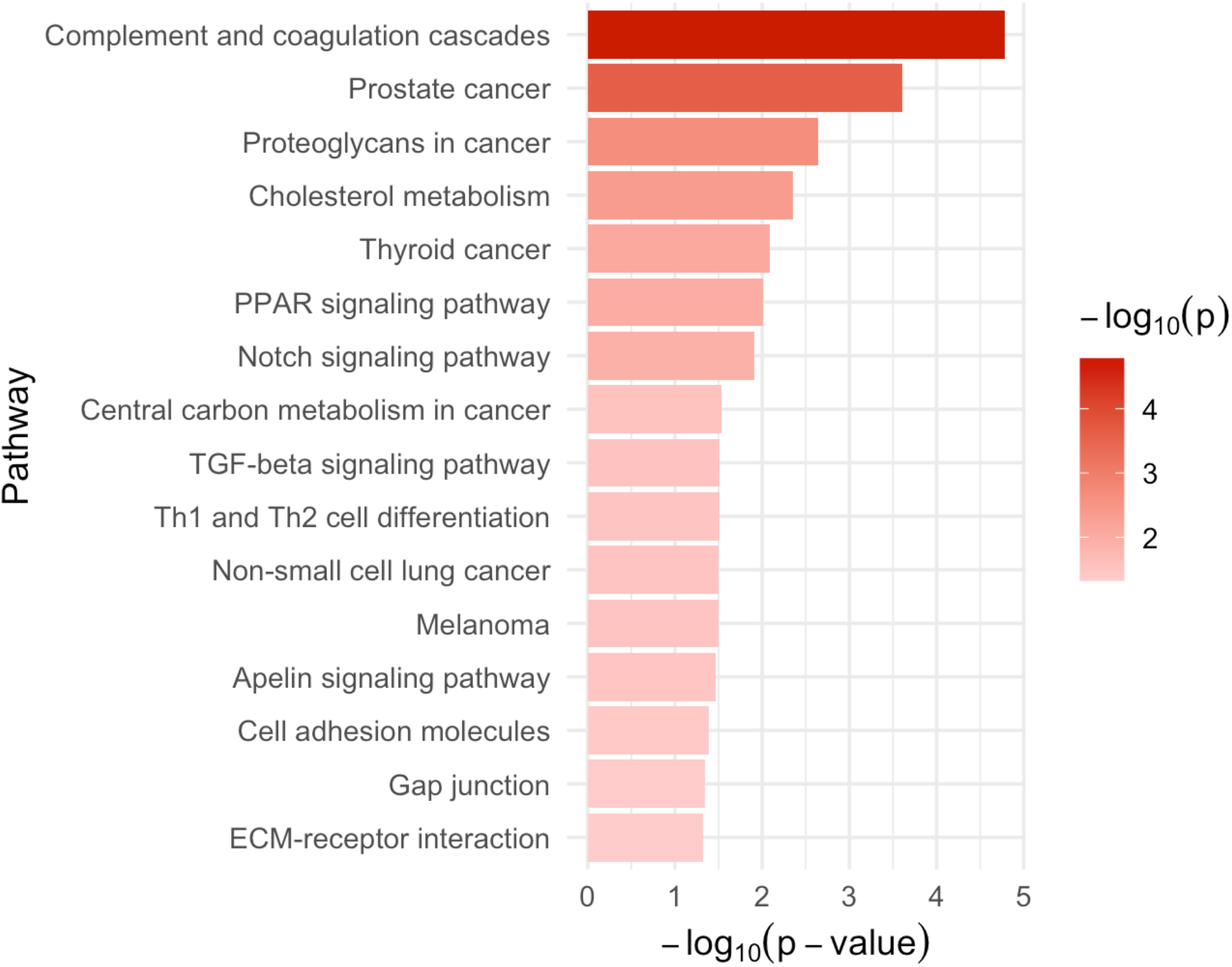
Pathways enriched in the set 44 proteins associated with hormonal contraceptive use, ranked by decreasing p-value of enrichment.

Several pathways related to inflammation and cellular communication such as Th1 and Th2 cell differentiation, ECM–receptor interaction, gap junctions, and cell adhesion molecules, were also enriched. Notably, a number of cancer-associated pathways (thyroid cancer, melanoma, non-small cell lung cancer) reached significance, potentially reflecting shared molecular mediators of cell proliferation and vascular remodeling. The list of significantly enriched pathways, including the adjusted p-values and the proteins associated with oral contraceptive use that contribute to each pathway, is provided in Supplemental Table T1.

### Gene Ontology Enrichment Analysis

Gene Ontology enrichment analysis revealed 25 GO terms significantly enriched biological processes and molecular functions (**Figure 4**) involving 39 of the 44 proteins associated with OC use. Notably, enriched categories include peptidase regulator activity, endopeptidase inhibitor activity, and serine-type endopeptidase inhibitor activity, which prominently feature proteins like SERPINA6, SERPINA7, FETUB, SPINK1, and PZP, all of which were elevated in OC users. These categories highlight functional roles related to hormone transport, protease regulation, and extracellular matrix remodeling.

**Figure 4:**
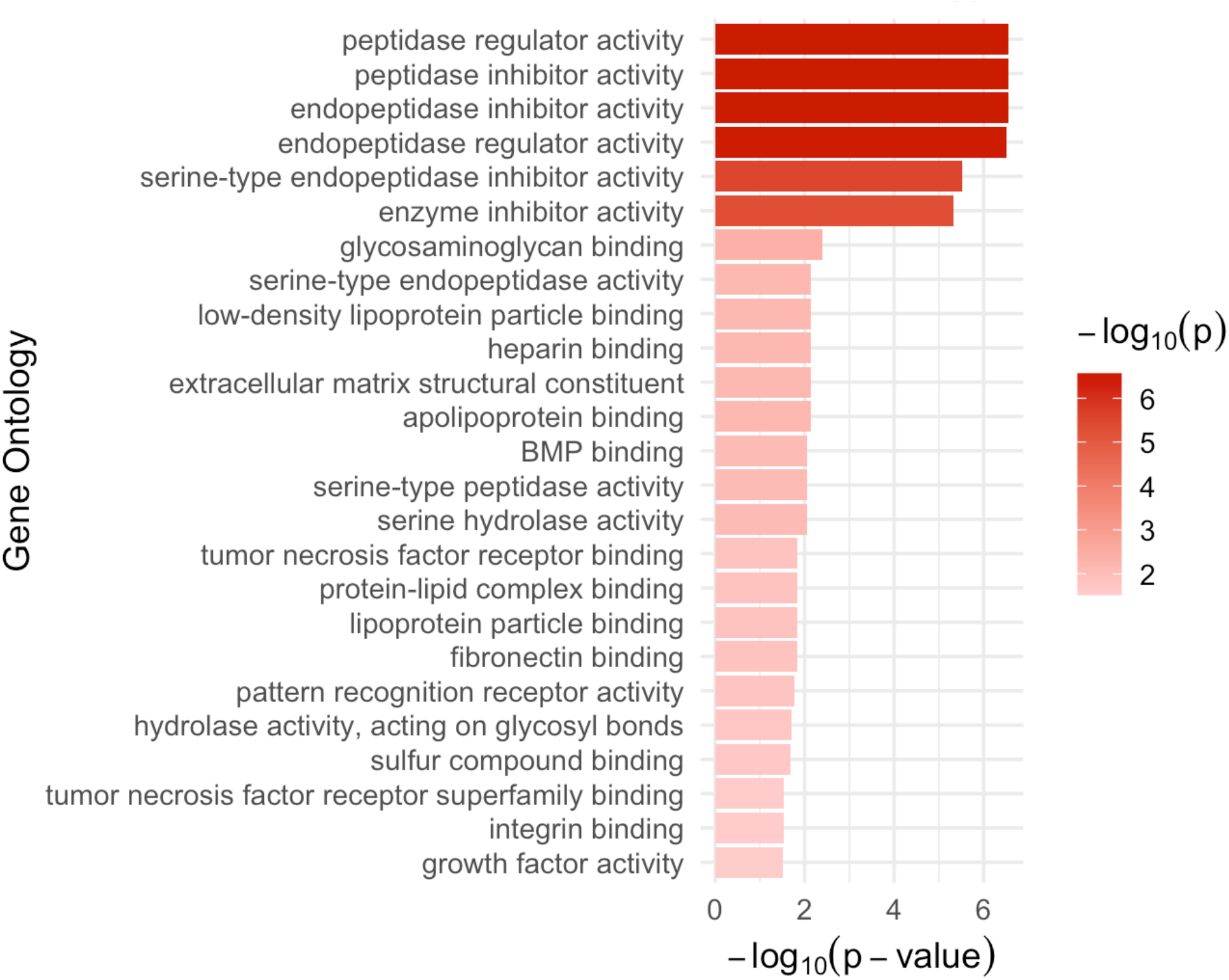
Gene Ontology terms enriched in the set 44 proteins associated with hormonal contraceptive use, ranked by decreasing p-value of enrichment.

Other enriched GO terms include complement activation and humoral immune response, with involvement of proteins such as PLG, ITIH3, and SERPINA6, pointing to immune-modulatory processes that may intersect with vascular and coagulation pathways. Additionally, categories related to lipoprotein particle remodeling and cholesterol efflux were also enriched, involving PLTP, ANGPTL3, and LPA, decreased in OC users.

The full list of enriched GO terms, adjusted p-values, and contributing proteins is provided in Supplemental Table T2, which includes all significantly enriched pathways and the subset of OC-associated proteins linked to each.

## DISCUSSIONS

The aim of this study was to identify proteomic alterations associated with oral contraceptive use in premenopausal African American women with stage 2 hypertension. Three key findings emerged. First, we identified 44 proteins significantly associated with OC use, encompassing both previously reported markers and novel candidates. Second, our results provide mechanistic support for established pathways of OC-induced hypertension, including activation of RAAS, heightened sympathetic nervous system activity, and altered regulation of antidiuretic hormone. Third, we highlight specific proteins such as SERPINA6, SERPINA7, and TNFSF13, that offer new insights into how OCs may influence cardiovascular risk in hypertensive women. Collectively, these findings suggest that OC use contributes to hypertension through multiple, interconnected molecular pathways.

One of the most important findings was the identification of proteins involved in the RAAS pathway, such as SERPINA6 (corticosteroid-binding globulin) and SERPINA7 (thyroxine-binding globulin), which were elevated in OC users. These proteins play important roles in hormone transport and may influence RAAS components, contributing to increased sodium and water retention and elevated BP ^11,12.^ Gene Ontology analysis further supports their involvement, highlighting enrichment in peptidase inhibitor activity, particularly among serpin family proteins which regulate protease activity essential for vascular stability and inflammatory control ^13,14.^ Additionally, TNFSF13 and TNFSF10, both associated with chronic inflammation, are consistent with the hypothesis that impaired RAAS feedback inhibition exacerbates hypertension in OC users ^15,16.^ These findings provide new insights into the inflammatory mechanisms linking OC use to BP regulation.

We also observed increased levels of SCG3 (secretogranin III), PLAU (plasminogen activator urokinase), and PLAUR (plasminogen activator urokinase receptor), supporting the hypothesis that OCs enhance sympathetic nervous system activity, which contributes to elevated BP. Pathway enrichment identified PLAU and PLAUR as part of the complement and coagulation cascades, suggesting that these proteins may play dual roles in sympathetic signaling and promoting a prothrombotic state in OC users ^17,18^. PLAU and PLAUR are expressed in sympathetic neurons and amplify neurogenic vascular responses and neurotransmitter release ^19-21^, thereby influencing vascular tone and BP regulation independently of plasmin generation. Although SCG3 is less directly studied in this context, related granin proteins such as chromogranin A are established modulators of catecholamine secretion and sympathetic outflow, suggesting a plausible role for SCG3 in OC-associated neuroendocrine activation ^22,23^. Collectively, these proteins are involved in neuroendocrine secretion, tissue remodeling, and inflammation, processes central to sympathetic tone and vascular function.

Furthermore, the changes in LPA and COL6A3 levels suggest that OCs may alter ADH regulation, thereby influencing fluid balance and extracellular matrix integrity, both important components of BP homeostasis. In parallel, LPA, along with PLTP and ANGPTL3, showed patterns consistent with OC-induced dyslipidemia, which may further impact vascular tone and inflammatory status ^24^. LPA induces vasoconstriction and stimulates intracellular calcium signaling in vascular smooth muscle cells, mechanisms known to trigger ADH release under osmotic and volume stress conditions ^25^. COL6A3, a structural ECM protein expressed in the kidney interstitium, is influenced by hormonal status and may alter renal responsiveness to ADH by modulating water reabsorption capacity and tubular architecture ^26-28^. Notably, ANGPTL3, a regulator of lipid metabolism, was decreased in OC users, suggesting complex interactions between OCs, lipid regulation, and hypertension ^29,30.^ In addition, NOTCH3, a receptor involved in vascular smooth muscle cell fate and remodeling, was differentially expressed and may act downstream of inflammatory and lipid-related signaling to influence arterial pressure regulation and vascular adaptation to hormonal exposure ^31^.

Several of the proteins identified, including PZP, SERPINA6, FETUB, PLG, PGLYRP2, and ITIH3, have been reported previously in large-scale population-based proteomic studies, such as the CHRIS and NESDA cohorts ^32,33.^ The consistency in direction and magnitude of associations for proteins such as PZP, SERPINA6 (linked to RAAS activation), FETUB, PLG (coagulation), PGLYRP2, and ITIH3 adds confidence to our findings. At the same time, we uncovered novel protein associations not previously linked to OC use. Notably, SMAD5, a receptor-regulated SMAD in the TGF-β/BMP signaling axis, emerged as a new candidate ^34^. SMAD5 mediates endothelial–mesenchymal transition and promotes extracellular matrix deposition and vascular smooth muscle proliferation, processes that are key to vascular remodeling and fibrosis in hypertension models ^35,36.^ These results suggest that hormonal modulation of the TGF-β/BMP–SMAD5 signaling axis may represent one mechanism through which OCs contribute to vascular remodeling and elevated blood pressure ^8^.

Taken together, these findings may help inform future efforts in CVD-related biomarker discovery and personalized contraceptive counseling for women with hypertension ^37^. A precision medicine approach, informed by proteomic profiling, could enable safer contraceptive choices that minimize cardiovascular risk in high-risk populations ^37^.

Our study has limitations. We did not collect data on OC formulation, dose, or duration, which limits assessment of specific OC types. The cross-sectional design precludes causal inference. Although we adjusted for important covariates, residual confounding cannot be ruled out. Finally, while most participants used pill-based OCs, potential misclassification of other hormonal contraceptive use may have diluted our estimates.

In conclusion, OC use in African American women with stage 2 hypertension is associated with proteomic changes across multiple pathways related to blood pressure regulation. Our findings confirm previously reported protein alterations while also identifying new candidate biomarkers and mechanisms that merit further investigation. This work underscores the importance of considering molecular heterogeneity in contraceptive risk and supports more individualized approaches to reproductive and cardiovascular care.

## ACKNOWLEDGEMENTS

This work was supported by the Chan Zuckerberg Initiative’s Foundation for Accelerate Precision Health Program to Advance Genomics Research at Meharry Medical College (CZIF2022-007043); by the National Institute of Health under award numbers EY033264, GM151274, UC2MD019626; and by the Veterans Affairs Office of Research and Development under award number CX002780. Dr Ayo Priscille Doumatey, National Institutes of Health – National Human Genome Research Institutes.

## Notes

**Conflict of Interest** The authors report no conflict of interest.

### Competing Interest Statement

The authors have declared no competing interest.

### Summary of Updates

A complete reinterpretation of the results, a revised framing of the hypothesis, and a significant refocusing of the introduction on hypertension. The following improvements have been made: - Rewritten the introduction to remain tightly focused on hypertension, removing tangential elements and better grounding the study in the context of hypertensive disorders in reproductive-age women. - Revised the hypothesis statement for clarity and fully restructured the results and discussion sections to reflect the new interpretation of findings. - Clarified the derivation of the analytic sample, providing a transparent account of how we arrived at 51 participants from an initial cohort of 669. This is now clearly outlined in the text and visually represented in a new figure.

